# Safe linkage of cohort and population-based register data in a genome-wide association study on health care expenditure

**DOI:** 10.1101/2020.10.17.334896

**Authors:** Eveline L. de Zeeuw, Lykle Voort, Ruurd Schoonhoven, Michel G. Nivard, Thomas Emery, Jouke-Jan Hottenga, Gonneke A.H.M. Willemsen, Pearl A. Dykstra, Narges Zarrabi, John D. Kartopawiro, Dorret I. Boomsma

**Affiliations:** Department of Biological Psychology, Vrije Universiteit, Amsterdam, the Netherlands; Amsterdam Public Health Research Institute, Amsterdam, the Netherlands; SURFsara, SURF, Amsterdam, the Netherlands; Statistics Netherlands, Den Haag/Heerlen, the Netherlands; Netherlands Interdisciplinary Demographic Institute, Den Haag, the Netherlands; Department of Public Administration and Sociology, Erasmus University, Rotterdam, the Netherlands; Amsterdam Reproduction and Development Research Institute, Amsterdam, the Netherlands

**Keywords:** Record linkage, Register-based data, Safely interconnected data, Genome-wide association study, Health care expenditure

## Abstract

**Background:** There are research questions whose answers require record linkage of multiple databases which may be characterized by limited options for full data sharing. For this purpose, the Open Data Infrastructure for Social Science and Economic Innovations (ODISSEI) consortium has supported the development of the ODISSEI Secure Supercomputer (OSSC) platform that allows researchers to link cohort data to data from Statistics Netherlands and run analyses in a high performance computing (HPC) environment.

**Methods:** After successful record linkage genome-wide association (GWA) analyses were carried out on expenditure for total health, mental health, primary and hospital care and medication. Record linkage for genotype data from 16,726 participants from the Netherlands Twin Register (NTR) with data from Statistics Netherlands was accomplished in the secure OSSC platform, followed by gene-based tests and estimation of total and SNP-based heritability.

**Results:** The total heritability of expenditure ranged between 29.4 (SE 0.8) and 37.5 (SE 0.8) per cent, but GWA analyses did not identify single SNPs or genes that were genome-wide significantly associated with health care expenditure. SNP-based heritability was between 0.0 (SE 3.5) and 5.4 (SE 4.0) per cent and was different from zero for mental health care and primary care expenditure.

**Conclusions:** We successfully linked genotype data to administrative health care expenditure data from Statistics Netherlands and performed a series of analyses on health care expenditure. The OSSC platform offers secure possibilities for analysing linked data in large-scale and realizing sample sizes required for GWA studies, providing invaluable opportunities to answer many new research questions.

**Key messages:**

- Cohort data of the Netherlands Twin Register were safely linked to population-based register data of Statistics Netherlands
- On the ODISSEI Secure Supercomputer (OSSC) platform genome-wide association analyses were carried out on linked genotype and health care expenditure data
- Variation in health care expenditure was for approximately one third explained by family-based heritability, but SNP-heritability based on genetic similarity across unrelated individuals explained only a very small proportion of variance
- The newly developed platform can serve as a prototype for realizing genome-wide association studies with sensitive data

## Introduction

Data collected for administrative or policy purposes often include detailed information at the individual level and are proving to be of great value for medical and scientific research. The Nordic countries are well-known for studies on health outcomes based on register data because these data can be linked across registers with a unique personal identification number^1^. Individuals’ data on for example, education or income, can be linked to the the use of social insurance schemes and health care, and also to data on their family members and their place of residence. The advantage of register data is that non-response is not a problem and that data are collected in a uniform manner from all individuals. However, register data are generally not collected with a specific research purpose in mind. Cohort studies on the other hand employ surveys, collect biological samples and carry out clinical and experimental studies to gather data directly from participants (lifestyle, personality, attitudes, genotypes, biomarkers, and clinical diagnoses). Linking individual-level register data to cohort data offers the possibility to enhance existing data resources and address research questions that are of relevance to individuals and to society.

In the Netherlands, population-based register data are collected and analysed by Statistics Netherlands (CBS; www.cbs.nl/en-gb) to publish reliable statistics on the Dutch economy and society (www.statline.nl). CBS has administrative data for the approximately 17 million inhabitants of the Netherlands on pedigree structure and outcomes, including education, income and health, that can all be linked at the individual level. CBS is legally entitled to make these data, under strict terms, available for research purposes as well as link them to external data in a secure remote-access (RA) environment (www.cbs.nl/microdata). However, this RA environment does not offer high-performance computing facilities which limits the scale of research projects. The Open Data Infrastructure for Social Science and Economic Innovations (ODISSEI; www.odissei-data.nl) has set up a sustainable national data infrastructure for research in the Netherlands by coordinating the integration of data from large cohort studies in the Netherlands and administrative data. ODISSEI has supported the development of a secure platform, called the ODISSEI Secure Supercomputer (OSSC)^2^ on the high-performance compute facility at *Samenwerkende Universitaire RekenFaciliteiten* (SURF; www.surf.nl) that houses the Dutch national supercomputer Cartesius with 2000 multi-core compute nodes, connected with Infiniband for high-speed, low-latency communication and storage access. The platform is based on customizable virtualized private clusters that are deployed on Cartesius, imposing strict security measures required by data owners and facilitates access and record linkage (see Figure S1).

In the current study we used the OSSC platform to link CBS register data on health care expenditure to genotype information from the Netherlands Twin Register (NTR; www.tweelingenregister.vu.nl). The NTR is a large twin-family study that has followed thousands of family members longitudinally with surveys and has collected DNA samples from a large number of their participants^3^. The scientific aim of our study was to run a genome-wide association (GWA) study and estimate the associations between genetic variants and health care expenditure. While the effect of environmental determinants related to overall health has been extensively studied, less is known about the genetic architecture of individual differences between people. Give that expenditure is directly related to medical conditions that are characterized by genetic contributions, we hypothesize that a substantial contribution of genetic differences to overall health expenditure exists. Knowledge on the genetic contributions to overall health is currently limited to studies that measured it with self-reports. Twin studies indicate that self-rated health is partly due to genetic differences between people with estimates of a heritability of over 30 per cent in young adulthood^4^ and almost 50 per cent at older ages^5^. The first GWA study (~6,700 individuals) on self-rated health did not report any genome-wide significant hits^6^, but a better powered study in almost 112,000 individuals identified 13 independent genome-wide significant signals, of which several were in regions previously implicated in specific diseases^7^. The proportion of variance in self-rated health explained by all the measured common genetic variants was 13 per cent. We argue that investigating overall health, more objectively measured by health care expenditure, will lead to a better understanding of its genetic architecture.

## Methods

### Participants

The Netherlands Twin Register (NTR) was established around 1987 by the Department of Biological Psychology at the Vrije Universiteit Amsterdam^3^. Adult twins, their families and parents of young twins take part in surveys. DNA collection from blood or buccal samples has been done in several large projects^3^. Genotyping has been carried out in subsamples that are, in general, unselected for specific traits. Height measured in centimetres was obtained from clinical and experimental studies or reported by participants in surveys. Many participants filled out more than one survey, and longitudinal height data were checked for consistency. NTR data collection was approved by the Central Ethics Committee on Research Involving Human Subjects of the VU University Medical Centre, Amsterdam, an Institutional Review Board certified by the U.S. Office of Human Research Protections (IRB number 00002991 under Federal-wide Assurance-FWA00017598; IRB/institute codes, NTR 03-180) and informed consent was obtained from all participants.

### Health care expenditure

Dutch residents are obliged by law (Health Insurance Act (ZVW)) to take out a basic health insurance. Merely 0.07 per cent of the population are conscious objectors and have no insurance for health care expenditure^8^. Statistic Netherlands (CBS) receives, via Vektis (www.vektis.nl), an executive organization of health insurance companies in the Netherlands, health care expenditure as reimbursed under the basic health insurance. Health care expenditure included the annual costs per resident for primary, hospital, mental health, birth, and geriatric care, physiotherapy and medication. Not included are costs that 1) fall under a supplementary health insurance, 2) fall outside of the Health Insurance Act and are paid by the patient or 3) fall under long-term care. We analysed average expenditure costs across all available reporting years (2009-2016) for total health, mental health, primary and hospital care and medication^9^. Data were log-transformed prior to analyses to correct for the skewness of the data.

### Genotype data

Genotyping was done on several platforms, i.e. Affymetrix-Perlegen, Illumina 660, Illumina Omni Express 1M, Affymetrix 6.0, Affimetrix Axiom and Illumina GSA. For criteria on quality control (QC) of the single nucleotide polymorphisms (SNP) and samples, before and after imputation, see^10^. Data were cross-platform phased and imputed using Mach-admix with GoNL^11^ as reference panel for all SNPs that were, after QC, present for at least one platform^12^. The cross-chip imputed dataset was used to calculate genetic principal components with the SmartPCA software^13^. Subsequently the dataset was aligned against the 1000G phase 3 version 5 reference panel and imputed on the Michigan imputation server^14^. Best guess genotypes were calculated for all SNPs in Plink 1.96^15^.

### Record linkage

Personal data (name, address, date of birth) of NTR participants and their Administrative number (A-number) under which Dutch residents are registered in the population register, are stored under a pseudonymized NTR identifier on a server that is disconnected from the internet (note that NTR does not have the social security number (BSN) of the participants)^16,17^. Phenotype data are stored under a different pseudonymized NTR identifier. The genotype data of NTR participants are stored at SURF in binary format and do not contain any identifiers, but the order within the dataset is an implicit identifier. At CBS all individual-level register data (microdata) are stored under a pseudonymized identifier (RIN) and CBS has access to the A-number of Dutch residents.

Figure 1 displays the different steps in the record linkage process. Record linkage between the NTR identifier and CBS identifier was done by the Central Record Linkage department (CBK) at CBS on the basis of the A-number (step 1). After linkage and removal of the A-number the NTR and CBS identifiers were encrypted and a record linkage key was sent to SURF (step 2). The key with encrypted identifiers was added to CBS microdata and NTR cohort data and the NTR cohort data were re-ordered (step 3). SURF, which acted as a trusted third party (TTP), used the record linkage key for pseudorandomizing the order of the genotype data (step 4). CBS uploaded CBS microdata and NTR cohort data to the OSSC environment via a secure virtual private network (VPN). SURF placed the re-ordered genotype data and the key of the new order of the genotype data in the OSSC (step 5). NTR linked CBS microdata to NTR cohort data and sorted the NTR cohort data in the same order as the new order of the genotype data (step 6). The complete procedure ensured that none of the parties involved had access to all the record linkage keys and therefore, could neither identify participants in the cohort nor in the administrative or genotype data.

**Figure 1.**
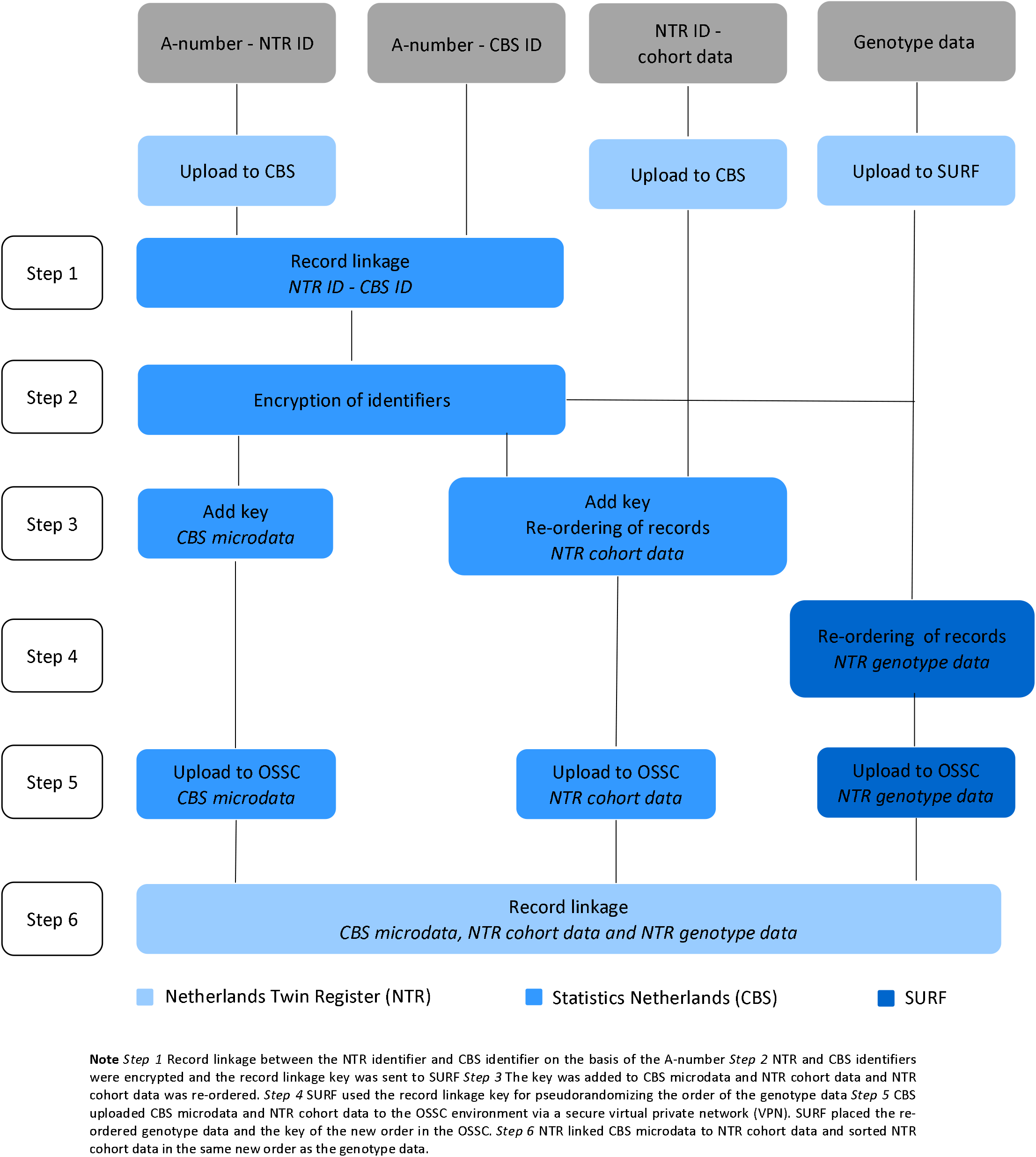
Flow chart of record linkage between data from the Netherlands Twin Register and data from Statistics Netherlands on the ODISSEI Secure SuperComputer (OSSC) platform.

### Statistical Analyses

To verify that the encryption of the NTR identifiers and the pseudo-randomization of the genotype data was correct, we conducted a GWA study on height reported by the NTR participants. The data on height were uploaded to the OSSC together with the other NTR cohort data, i.e. data on covariates, and underwent the same procedure of identifier encryption and the analyses were carried out using the re-ordered genotype data. A large meta-analysis reported genetic variants associated with height^18^ and we estimated the genetic correlation between these and our own results with LD-score regression^19^. For a successful procedure, we expected a genetic correlation close to one.

We regressed the log-transformed total health, mental health, primary and hospital care and medication expenditure on all SNPs in linear mixed models in GCTA 1.92.1beta6^20^, controlling for genetic relatedness by including a SNP-derived genetic relationship matrix (GRM) with all the off-diagonal elements < 0.05 set to zero. Population stratification was taken into account by including 10 principal components (PC) from the PCA of the SNP genotypes as fixed effects^13^. Sex, age and age squared also were included as fixed effects. To declare genome-wide significance for SNP-phenotype associations an alpha level of 5 × 10^-8^ was adopted^21^. Based on the GWA results we conducted genebased association analyses in MAGMA^22^. Genetic variants were assigned to genes on the basis of their position according to the NCBI 37.3 build, resulting in 15,438 genes. The European panel of the 1000 genomes^23^ data was used as a reference for linkage disequilibrium. In line with the Bonferroni method a genome-wide significance threshold of 0.05 / 15,438 = 3 × 10^-6^ was adopted for genebased association tests.

We estimated family-based heritability employing the differences in genetic resemblance between all family members in our sample as determined with the GRM in GCTA 1.92.1beta6^20^. The SNP-based heritability for each of the outcome measures and we estimated the genetic correlation with self-rated health^7^ were estimated with LD-score regression v1.0.0^19^.

## Results

Genotype data were successfully linked to CBS for 16,726 individuals from European ancestry^24^, who had given consent for linking to external databases and for whom the Administrative number was known. The analysis of adult height data (N = 12,498) gave a SNP-based heritability of 37.3 (5.9) per cent. The genetic correlation between our results and the results from the large meta-analysis on height as reported by the GIANT consortium was .99 (.06), indicating that the record linkage was done correctly.

For 14,572 participants (5,842 males and 8,727 females) with an average age of 45, from 5,669 families (average family size of 2.6) genotype data and expenditure for total health care in euros were available (males: mean = 1556.7, SD = 3406.9 females: mean = 1668.8, SD = 3432.1), mental health care (males: mean = 113.2, SD = 835.1; females: mean = 158.0, SD = 1308.7), medication (males: mean = 175.1, SD = 784.1; females: mean = 206.6, SD = 631.3), primary care (males: mean = 117.4, SD = 49.1; females: mean = 133.7, SD = 61.1) and hospital care (males: mean = 813.4, SD = 2292.9; females: mean = 1009.4, SD = 2260.9) (see Figure 2 and S2). The unavailability of health care expenditure data might have been due to, for example, residence outside of the Netherlands.

**Figure 2.**
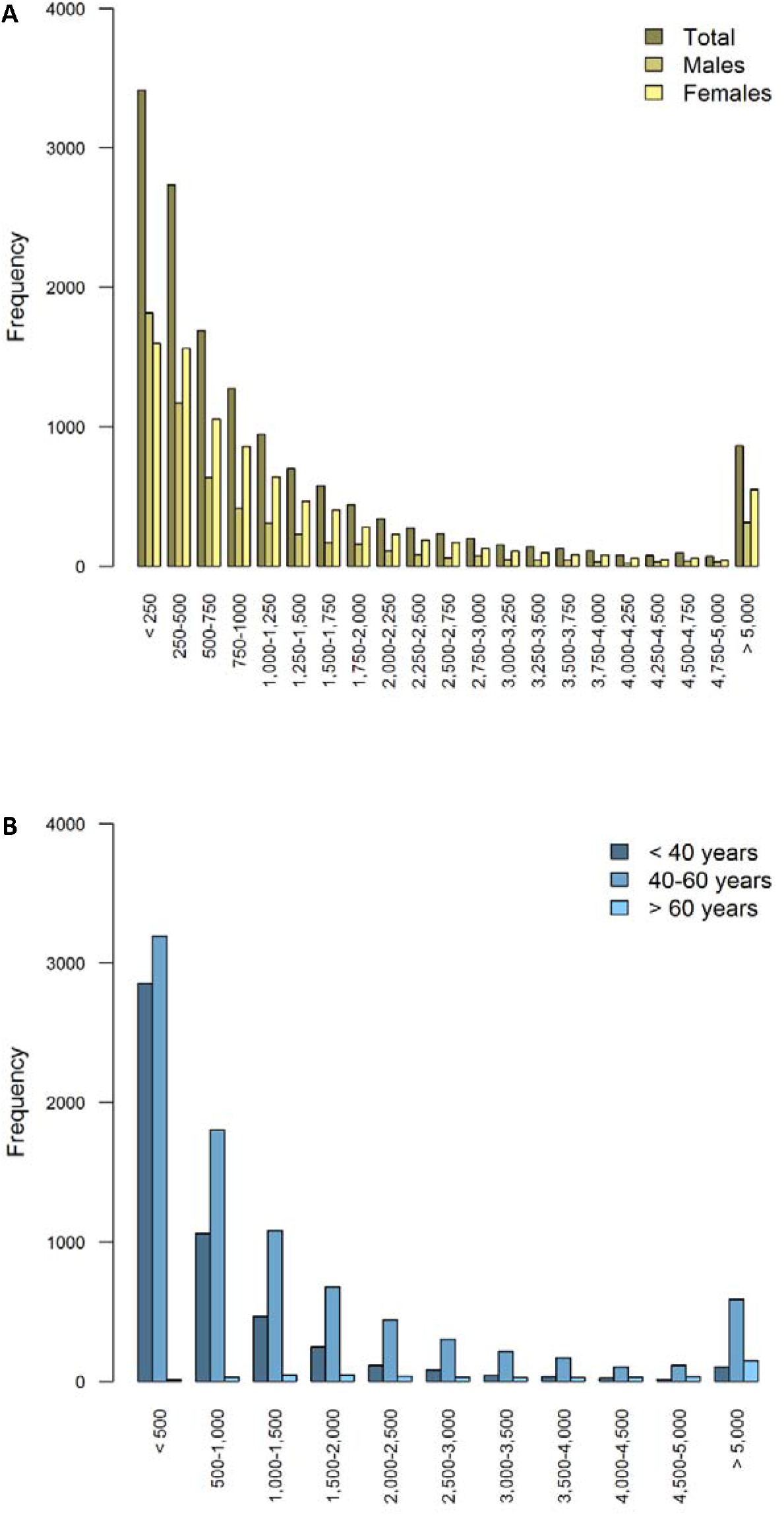
Histogram of total health care expenditure in Euros separately for (A) males (N = 5,845) and females (N = 8,727) and (B) for individuals < 40 years (N = 5,059), between 40 and 60 years (N = 8,706) and > 60 years (N = 483)

The results of the genome-wide association analyses summarized in Manhattan and QQ plots for health care expenditure are depicted in Figure 3 and S3-S6. There were no SNPs that were associated with health care expenditure at a genome-wide significance level and gene-based tests did not reveal genes that were significantly associated with health care expenditure with the strongest associations for the genes TRPV3 (17p13, *p* = 2 × 10^-5^), CAT (11p13, *p* = 7 × 10^-5^) and SSBP2 (5q14, *p* = 4 × 10^-5^). Family-based heritability was .319 (.01) for total health care, .302 (.01) for mental health care, .294 (.01) for medication, .375 (.01) for primary health care and .295 (.01) for hospital care and SNP-based heritability was −004 (.03) for total health care, .054 (.03) for mental health care, -.005 (.04) for medication, .044 (.04) for primary health care and -.002 (.04) for hospital care (see Figure 4). The genetic correlation between the GWA results that showed a significant SNP-based heritability and the results from the meta-analysis on self-rated health^7^ was .26 (.17) for mental health care expenditure and .70 (.30) for primary care expenditure.

**Figure 3.**
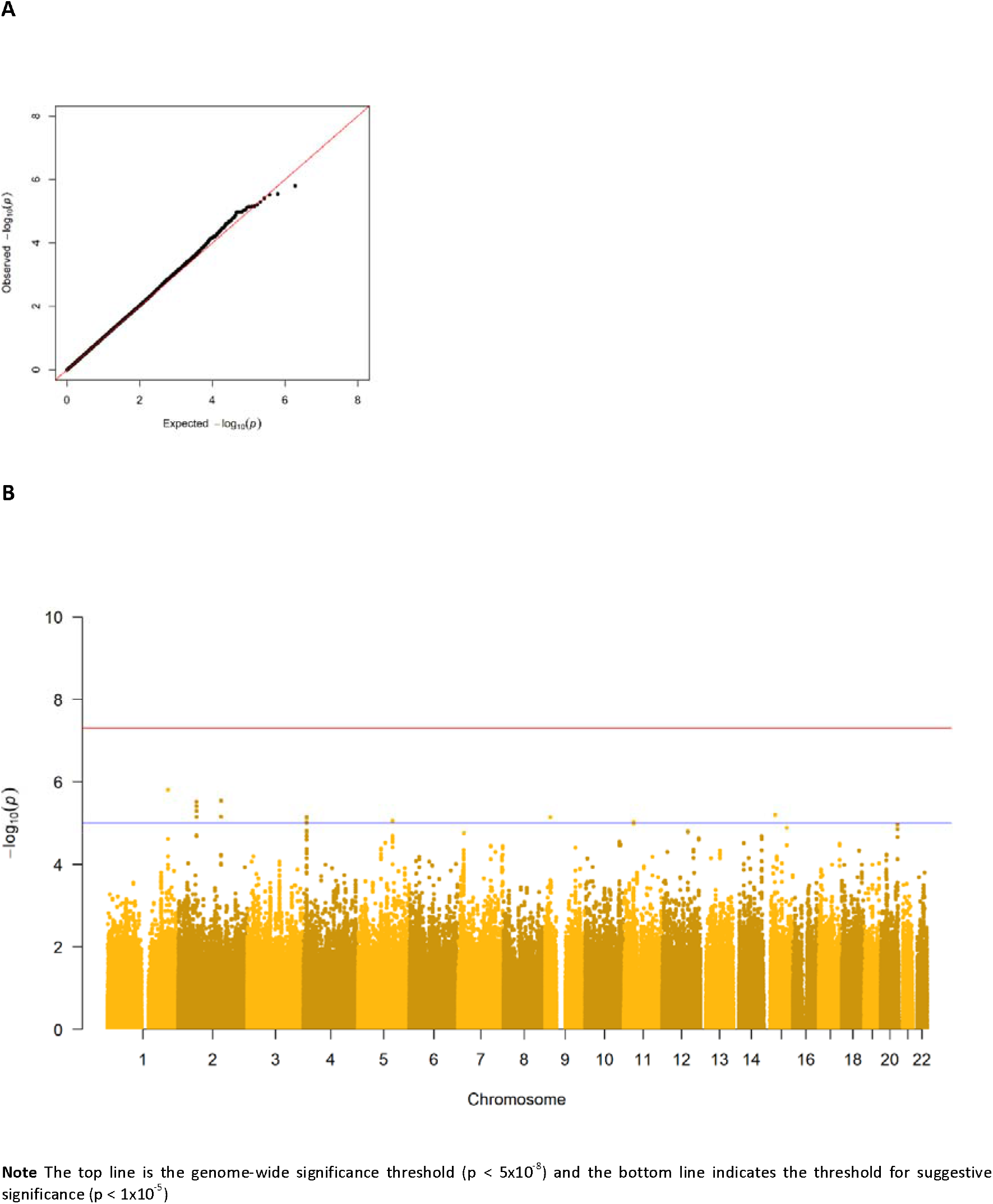
(A) Q-Q plot and (B) Manhattan plot of p-values of the genome-wide association analysis for total health care expenditure.

**Figure 4.**
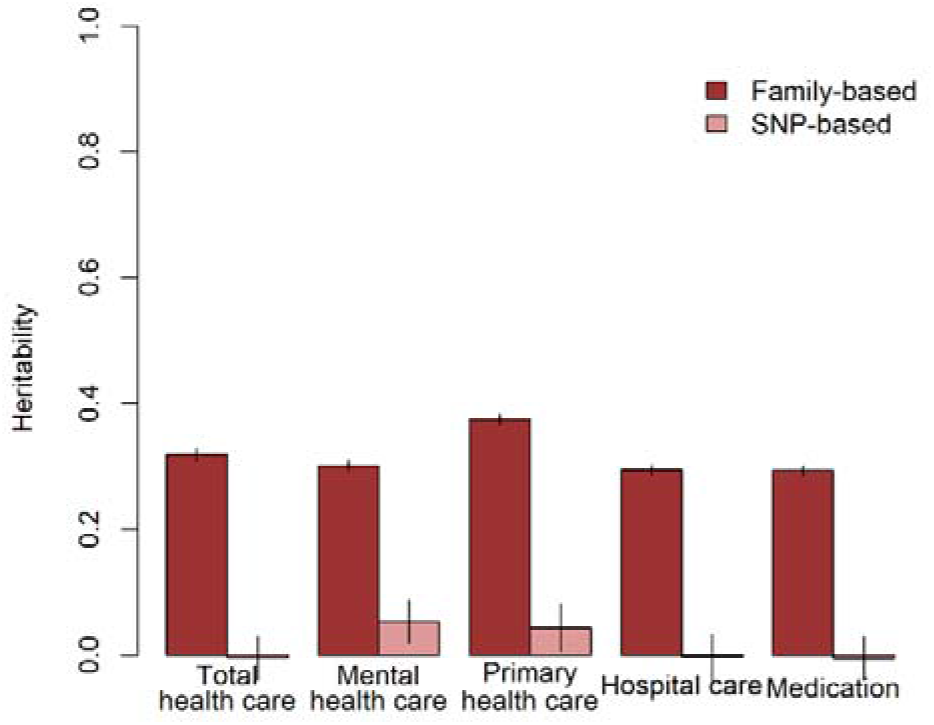
Family-based and SNP-based heritability of total health, mental health, primary health and hospital care and medication expenditure.

## Discussion

The aim of the current study was to demonstrate the feasibility of running a large-scale genome wide association (GWA) study for health care expenditure by linking NTR genotype data and CBS administrative data on the OSSC platform. We successfully linked genotype data to self-reported height from NTR participants in the OSSC after encrypting all identifiers and pseudo-randomization of the order in the genotype data. The finding of a significant SNP-heritability for height and a genetic correlation of unity with the GIANT study on height^18^ confirmed that the encryption and reordering of the data was done correctly. The OSSC thus enabled us to link sensitive data from multiple databases in a privacy protecting way on a secure system that provided the required high-performance computing facilities.

No genetic variants that were genome-wide significantly associated with health care expenditure were found. For significantly associated genes at the stringent genome wide level, larger studies are required, and this will become feasible when multiple Dutch cohorts collaborate in the OSSC environment. No significantly associated genes were found, but we identified some promising genes, e.g. SSBP2, that were previously found to be related with diseases^25^. Overall, the proportion of variance explained by all genotyped autosomal SNPs ranged from 0 to 5.4 per cent. The estimate for the SNP-based heritability was different from zero for mental health care expenditure and primary care expenditure. The genetic correlation between self-rated health and health care expenditure was larger for primary care (r = .66) compared to mental health care (r = .26). Family-based heritability was much larger for all types of health care expenditure (29-38 per cent), indicating that there are possibilities for larger samples to identify genetic variants related to health care expenditure. Health care expenditure may be influenced by a large number of genetic variants that all have a very small effect, but also by rare genetic variants that, although they can have a large effect, explain only a small part of the differences between people. Larger sample sizes are required to identify the effects of common genetic variants on health care expenditure. The OSSC brings these large sample sizes within reach as it will provide the possibility to link multiple cohorts to the register data allowing the exact same outcome measure to be analysed across all these cohorts.

Identifying genetic variants associated with overall health is important as these genetic variants can be employed in bidirectional Mendelian randomization (MR)^26^. MR is a natural experiment that employs genetic variants as instrumental variables and which has been suggested to provide the best opportunity to establish the causal effect of specific traits on overall health^27^, since randomized controlled trials are in this case impossible to implement. The identification of the impact of specific traits on overall health will give information needed to estimate the costeffectiveness of prevention and intervention programs. Such information is essential as health care expenditure in the Netherlands has increased to 76.9 billion euros, 9.9 per cent of the gross domestic product (GDP)^28^, only partly explained by the growth of the proportion of the Dutch population over 65 years^29^.

The OSSC development is moving towards a production-ready version of the platform by automating the manual steps for setting up the virtual environment and creating a scalable and robust solution. The platform allows data- and compute-intensive research projects to be conducted in parallel. The individual-level register data from Statistics Netherlands will offer opportunities for research in public health, health care, economics, education, sociology, bioinformatics, ‘omics’, and many other fields. The OSSC will also facilitate multidisciplinary research. Projects may for example, link data from MRI scans to data on psychiatric diagnoses as well as to data about urbanization level of the neighbourhood to get more insight into the mediating effect of neural processes in the association between the urban environment and psychopathology^30^. In addition, in light of the current Covid-19 pandemic the linkage of this population-based register data with cohort data will provide researchers with invaluable opportunities to determine the impact of the pandemic and the lockdown measures on individual outcomes such as mental health and educational outcome in children, who are currently being home schooled, or address questions on host-virus interactions.

## Acknowledgements

We are thankful to the twin families registered with the Netherlands Twin Register for their participation. We gratefully acknowledge Open Data Infrastructure for Social Science and Economic Innovations (ODISSEI) (NWO: NRGWI.obrug.2018.008)’; ‘Netherlands Twin Register Repository: researching the interplay between genome and environment’ (NWO: 480-15-001/674); ‘KNAW Academy Professor Award’ (PAH/6635); ‘Genetics as a research tool: A natural experiment to elucidate the causal effects of social mobility on health’ (ZonMw: 531003014). This work was carried out on the Dutch national e-infrastructure with the support of SURF Cooperative. The results are based on own analyses of the VU Amsterdam researchers based on the non-public data from Statistics Netherlands.

