## Supplements for "Safe linkage of cohort and population-based register data in a genome-wide association study on health care expenditure"

**Figure S1 *Architecture of the ODISSEI Secure Supercomputer (OSSC)***

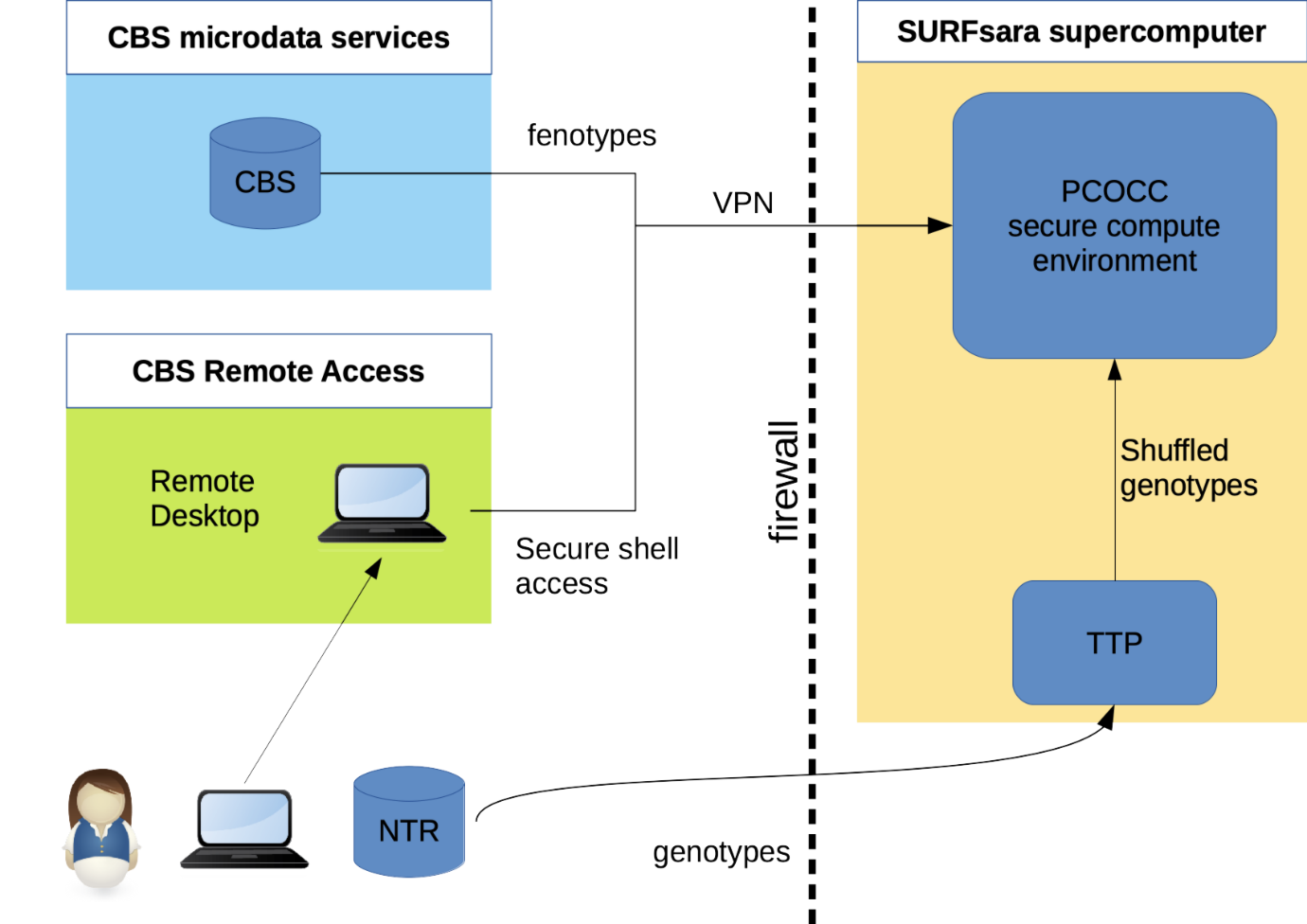

**Figure S2 *Histogram of health care expenditure for (A) mental health care, (B) medication, (C) primary care and (D) hospital care***

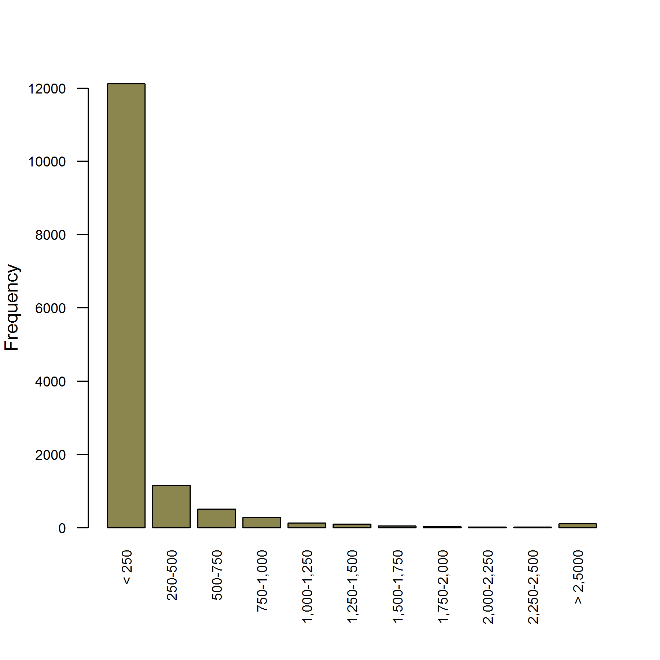

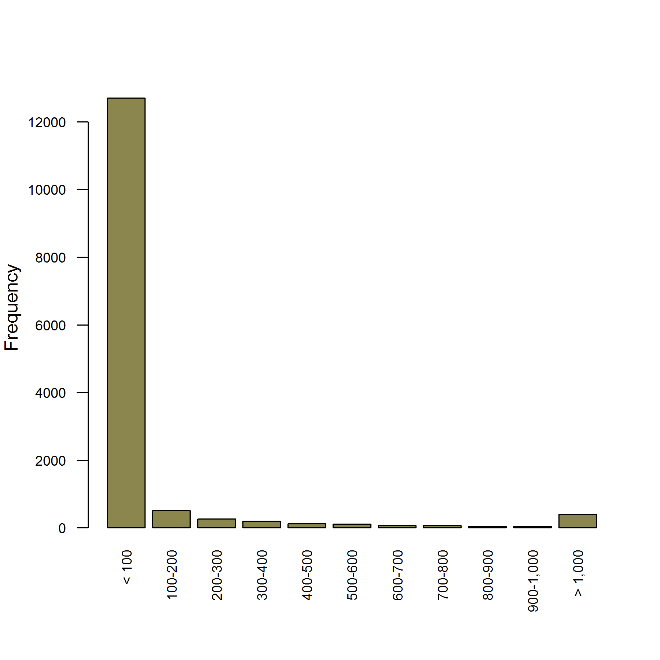

**A	B**

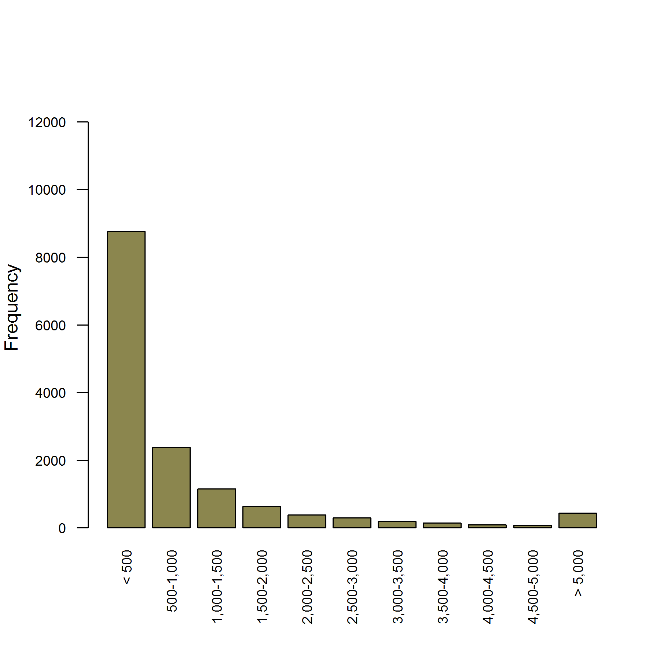

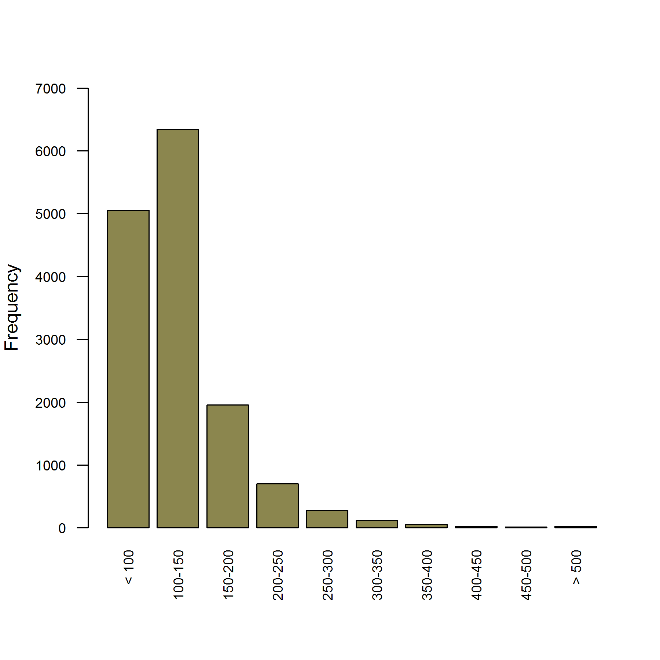

**C	D**

**Figure S3 *(A) Q-Q plot and (B) Manhattan plot of P-values of the genome-wide association analysis for mental health care expenditure***

**A**

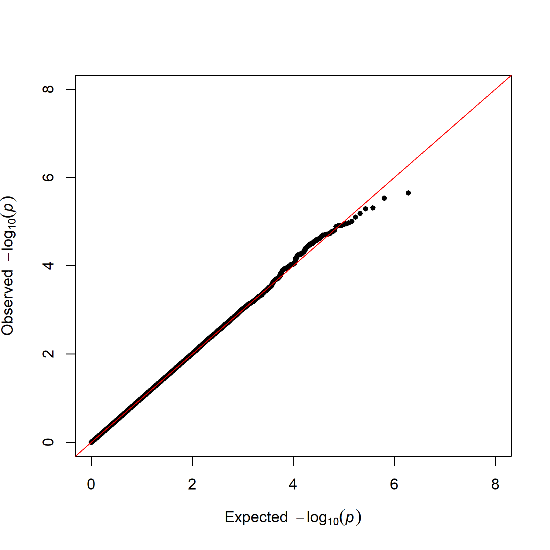

**B**

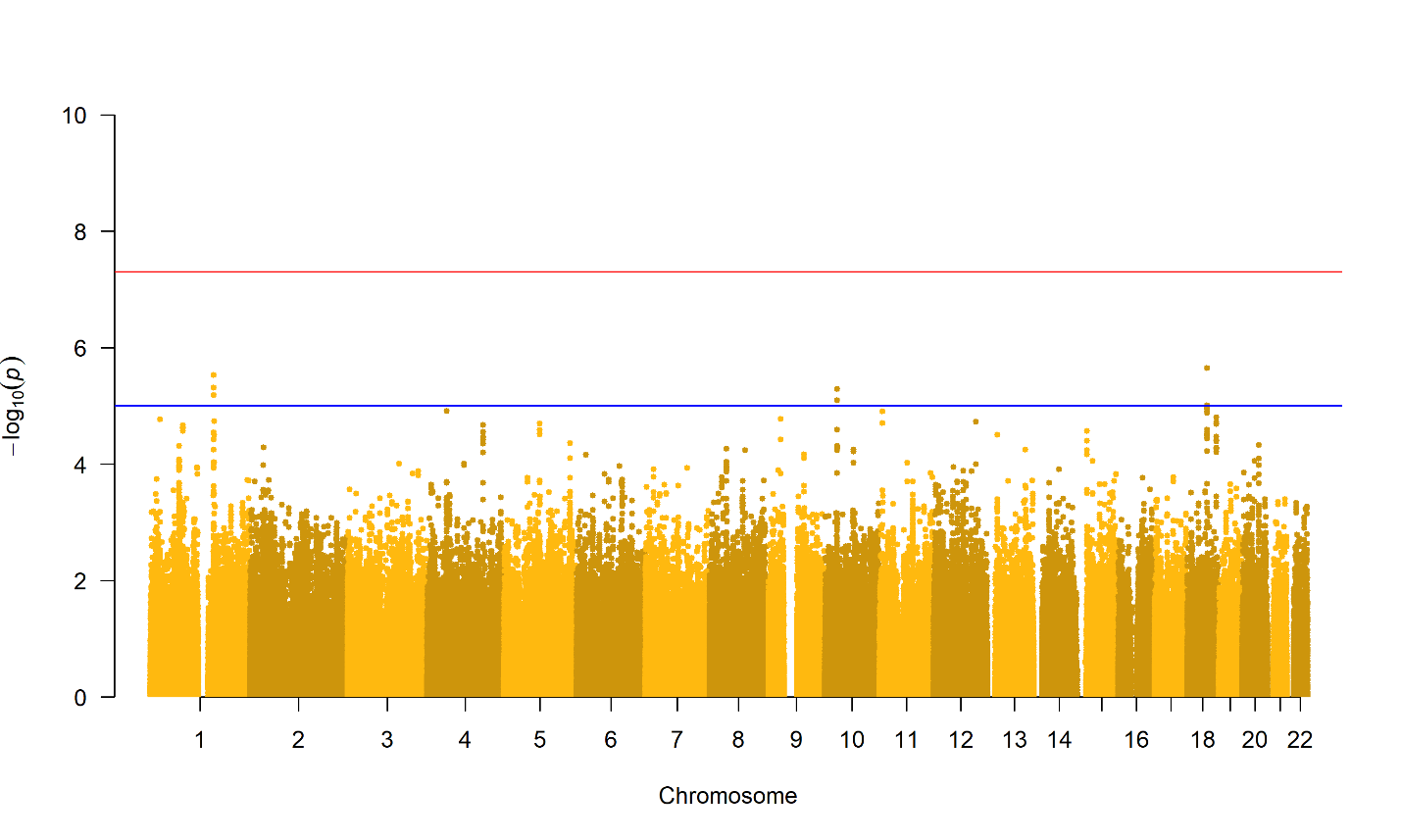
**Note** The top line is the genome-wide significance threshold (p < 5x10^-8^) and the bottom line indicates the threshold for suggestive significance (p < 1x10^-5^)

**Figure S4 *(A) Q-Q plot and (B) Manhattan plot of P-values of the genome-wide association analysis for medication expenditure***

**A**

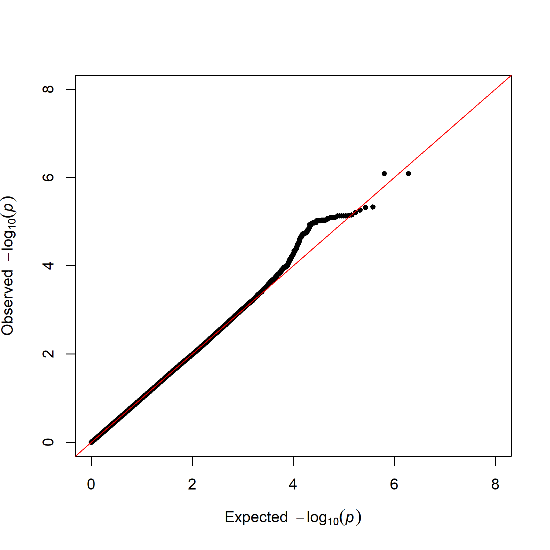

**B**

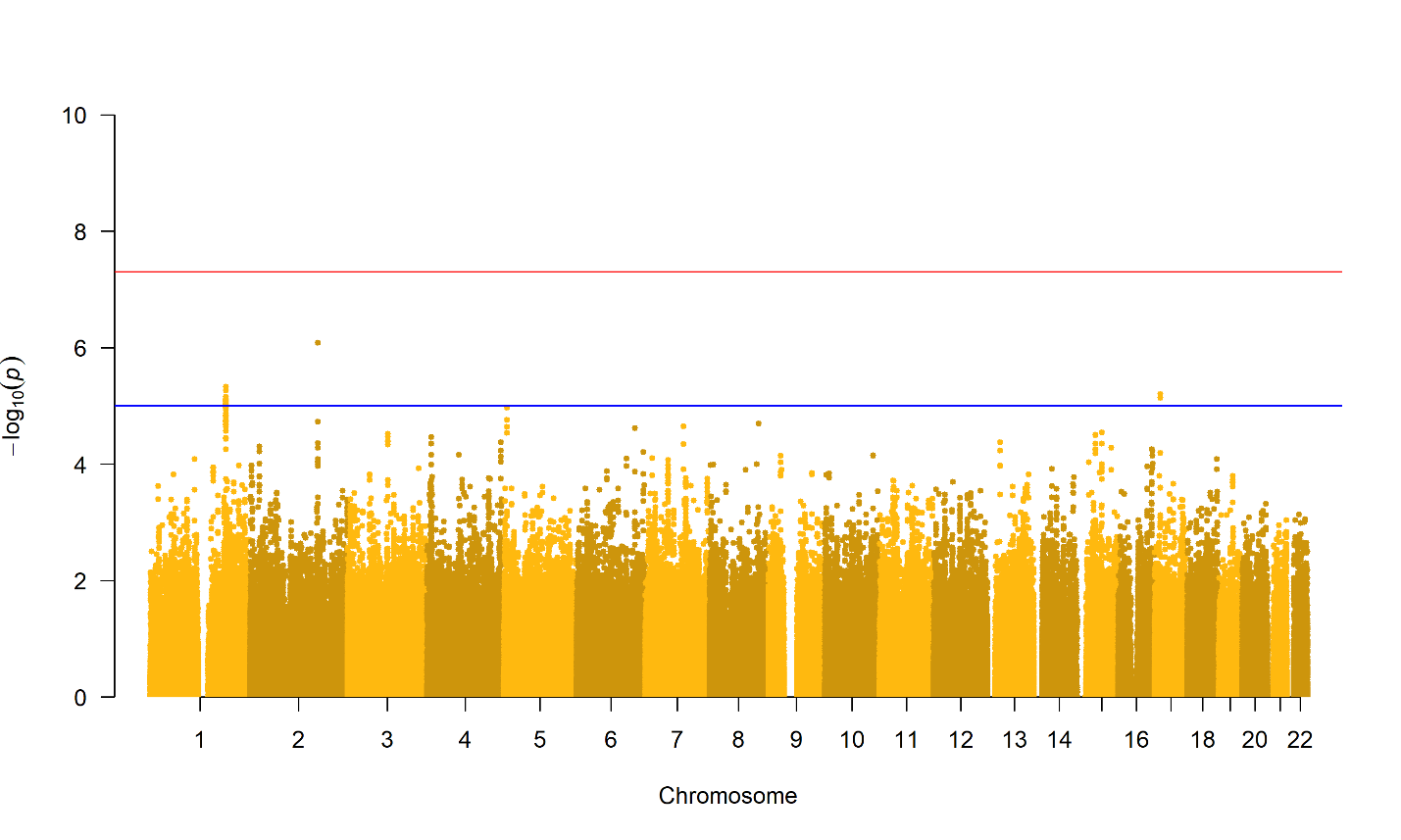

**Note** The top line is the genome-wide significance threshold (p < 5x10^-8^) and the bottom line indicates the threshold for suggestive significance (p < 1x10^-5^)

**Figure S5 *(A) Q-Q plot and (B) Manhattan plot of P-values of the genome-wide association analysis for primary care expenditure***

**A**

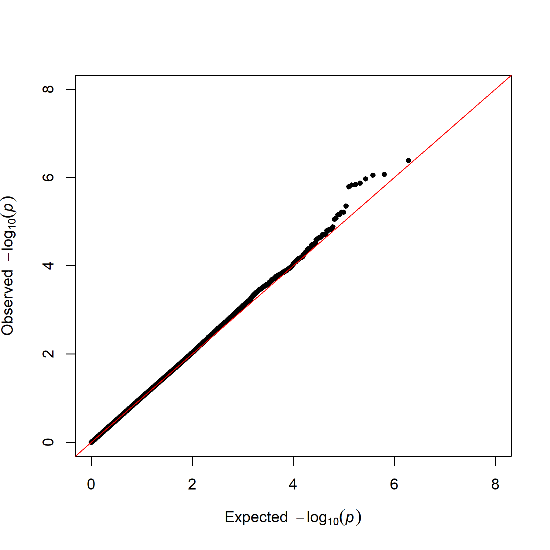

**B**

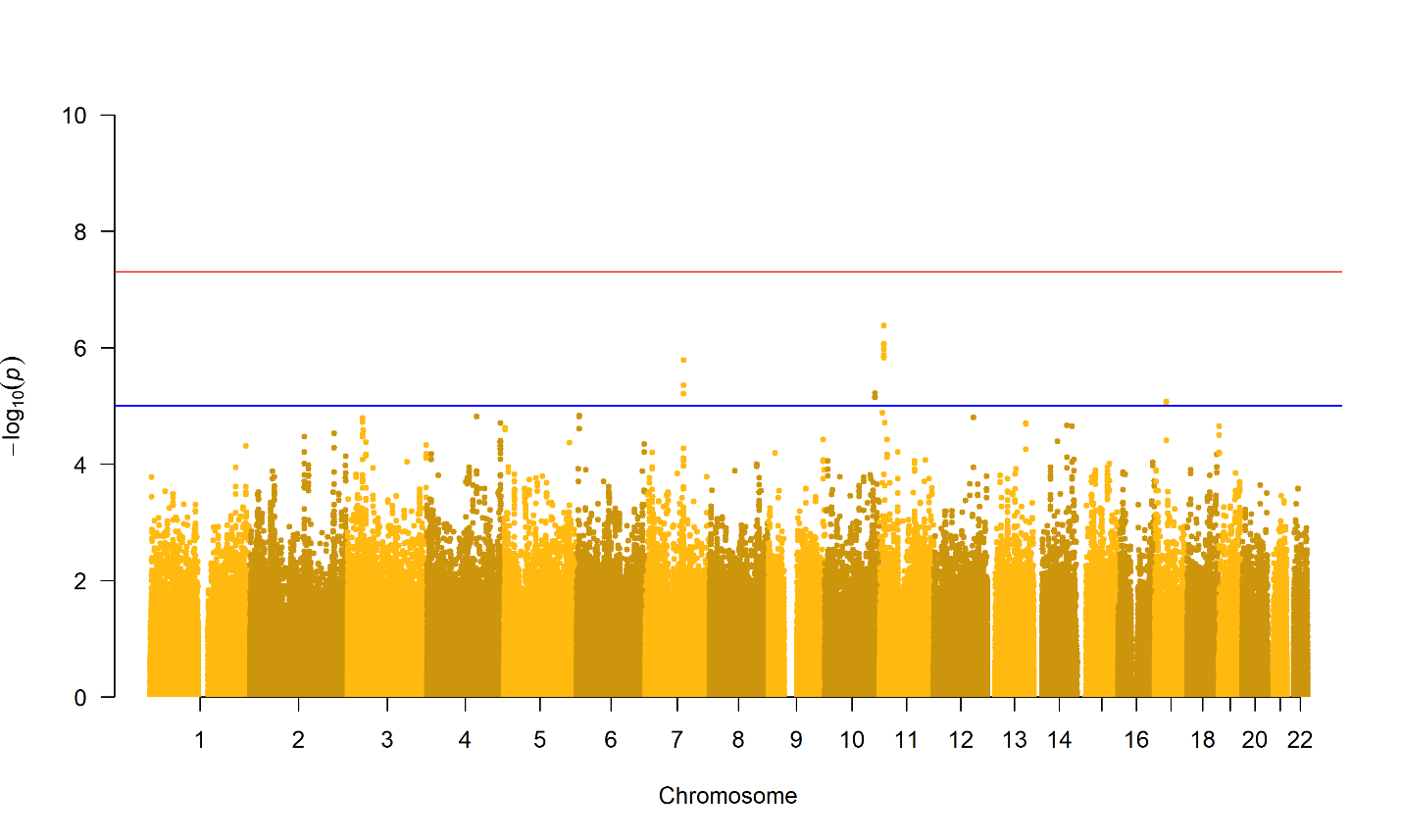

**Note** The top line is the genome-wide significance threshold (p < 5x10^-8^) and the bottom line indicates the threshold for suggestive significance (p < 1x10^-5^)

**Figure S6 (*A) Q-Q plot and (B) Manhattan plot of P-values of the genome-wide association analysis for hospital care expenditure***

**A**

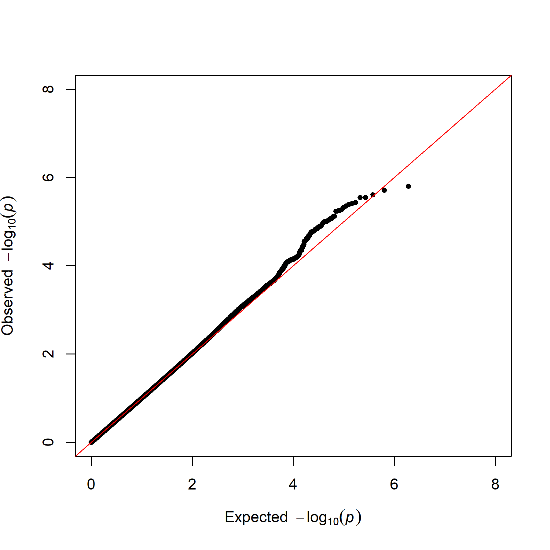

**B**

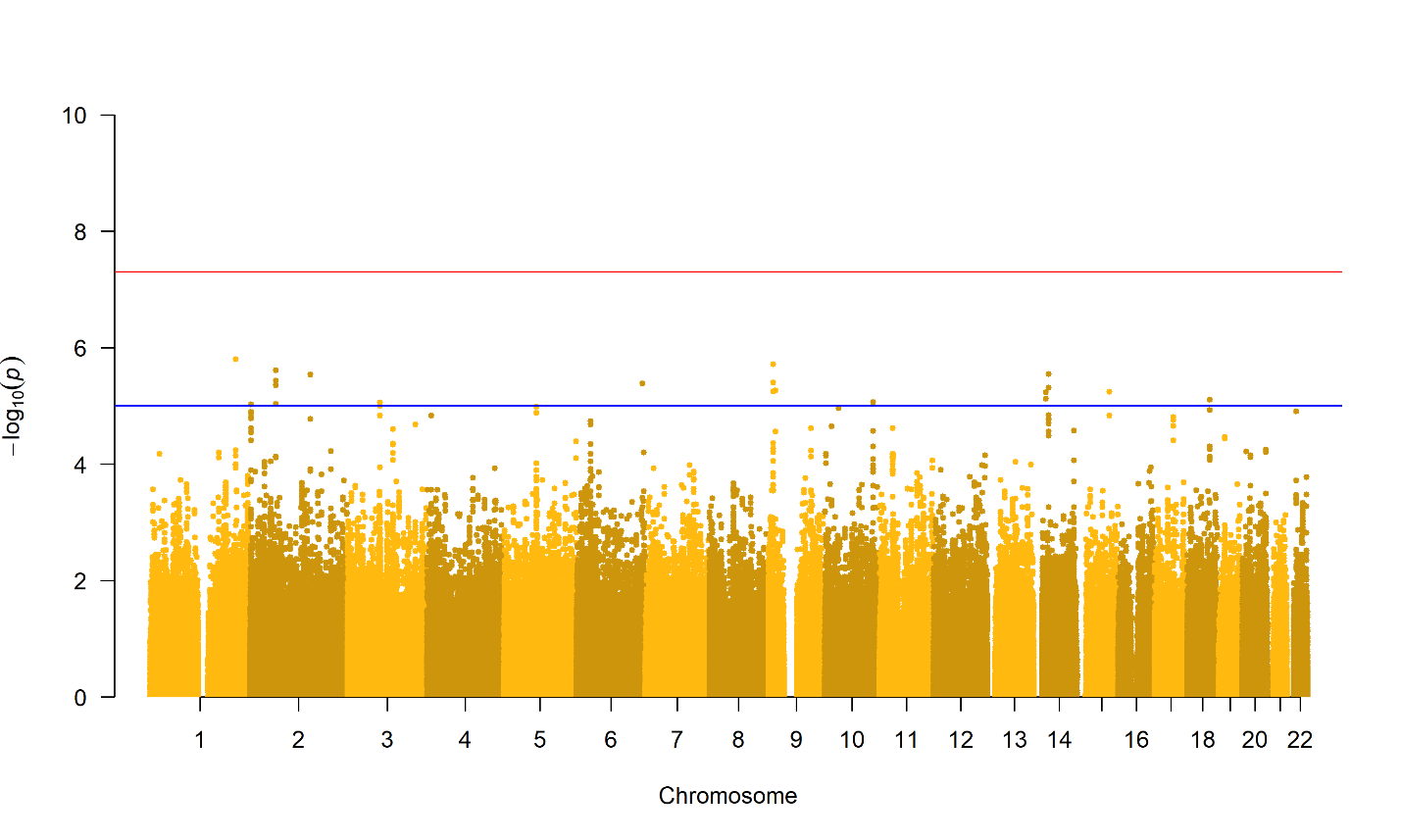

**Note** The top line is the genome-wide significance threshold (p < 5x10^-8^) and the bottom line indicates the threshold for suggestive significance (p < 1x10^-5^)
